# Pigs as a new behavioral model for studying Pavlovian eyeblink conditioning

**DOI:** 10.1101/2021.04.02.438144

**Authors:** Henk-Jan Boele, Sangyun Joung, Joanne E. Fil, Austin T. Mudd, Stephen A. Fleming, Sebastiaan K. E. Koekkoek, Ryan N. Dilger

## Abstract

**Intro:** Pigs have been an increasingly popular preclinical model in nutritional neuroscience, as their anatomy, physiology, and nutrition requirements are highly comparable to those of humans. Eyeblink conditioning is one of the most well-validated behavioral paradigms in neuroscience to study underlying mechanisms of learning and memory formation in the cerebellum. Eyeblink conditioning has been performed in many species but has never been done on young pigs. Therefore, our aim here was to develop and validate an eyeblink conditioning paradigm in young pigs.

**Method:** Eighteen intact male pigs were artificially reared from postnatal day 2 to 30. The eyeblink conditioning setup consisted of a sound-damping box with a hammock that pigs were placed in, which allowed the pig to remain comfortable yet maintain a typical range of head motion. In a delay conditioning paradigm, the conditional stimulus (CS) was a 550 ms blue light-emitting diode (LED), the unconditional stimulus (US) was a 50 ms eye air-puff, the CS-US interval was 500 ms. Starting at postnatal day 14, pigs were habituated for five days to the eyeblink conditioning setup, followed by 5 daily sessions of acquisition training (40 paired CS-US trials each day).

**Results:** The group-averaged amplitude of eyelid responses gradually increased over the course of the five days of training, indicating that pigs learned to make the association between the LED light CS and the air-puff US. A similar increase was found for the conditioned response (CR) probability: the group-averaged CR probability on session 1 was about 12% and reached a CR probability of 55% on day 5. The latency to CR peak time lacked a temporal preference in the first session, but clearly showed preference from the moment that animals started to show more CRs in session 2 and onwards whereby the eyelid was maximally closed exactly at the moment that the US would be delivered.

**Conclusion:** We concluded that 4-week-old pigs have the capability of performing in a cerebellar classical association learning task, demonstrating for the first time that eyeblink conditioning in young pigs has the potential to be a valuable behavioral tool to measure neurodevelopment.

## 1. Introduction

The use of pigs as an experimental animal model has been increasing in various fields, including neuroscience (Gieling et al., 2011; Kornum & Knudsen, 2011; Lind et al., 2007) and pediatric nutrition (Fleming et al., 2019; Liu et al., 2014; Rytych et al., 2012), for its several major advantages. First, pigs share gross neuroanatomical similarities to humans (Dickerson & Dobbing, 1967). Second, pigs are precocial in nature, which allows them to be weaned at birth, raised in controlled environments, and trained on behavioral paradigms early in life to assess various cognitive capacities, such as sensory discrimination, spatial learning and memory, and recognition memory (Friess et al., 2007; Wang et al., 2007; Fleming & Dilger, 2017). Third, young pigs have similar gastrointestinal anatomy, physiology, and nutrient requirements to infants, which makes them a great preclinical model for pediatric nutrition (Odle et al., 2014). For these reasons, the use of young pigs as a translational model in nutritional and developmental neuroscience is increasing (Mudd & Dilger, 2017).

In the field of nutritional and developmental neuroscience, pigs have been tested in a variety of behavioral tasks, including T- maze, radial arm maze, and novel object recognition (Bolhuis et al., 2004; Dilger & Johnson, 2010; Fleming & Dilger, 2017). Most of these tasks focus on the function of the hippocampus and/or the neocortex. However, pigs have not been extensively investigated in a task that focuses specifically on the cerebellar motor response in eyeblink conditioning. Therefore, our aim was to establish procedures to conduct Pavlovian eyeblink conditioning in pigs, which is a cerebellar-dependent learning task. During eyeblink conditioning, subjects typically hear a short beep or see a light flash (conditional stimulus, CS), followed several hundred milliseconds later by an air-puff on the eye (unconditional stimulus, US). In a cerebellar-dependent ‘delay paradigm’, the CS and US have different onset delays but co-terminate **(Figure 1A, B)**. As a result of repeated CS-US pairings, subjects eventually associate the CS with US, and in anticipation of the US learn to close their eyes in response to the CS. This anticipatory behavior to close the eye after the CS but before the US is called the conditioned response (CR) (Ten Brinke, 2013; Heck et al., 2013; Freeman & Steinmetz, 2011). Eyeblink conditioning became a popular learning model because it is simple in its form but is discrete in that it specifically measures associative and sensory-motor learning (Heiney et al., 2014). As a result, the neural circuits and plasticity mechanisms in cerebellum involved in eyeblink conditioning have been studied in very high detail **(Figure 1C)**. Eyeblink conditioning has been performed in many species, including humans (Cason, 1922; Oristaglio et al., 2013; Thürling et al., 2015), cats (Woody & Brozek, 1969), ferrets (Svensson et al., 1997), rabbits (for instance: Gormezano et al., 1962; McCormick et al., 1982), rodents (for instance: Albergaria et al., 2018; Boele et al., 2010; Heiney et al., 2014), and sheep (Johnson et al., 2008). To our knowledge, eyeblink conditioning has never been performed on young pigs. Still, eyeblink conditioning may be a suitable behavioral paradigm for the pig because it can accurately measure sensitive changes in behavioral and cognitive development in early life, when other behavioral paradigms may not be applicable yet (Reeb-Sutherland & Fox, 2013). It can also be utilized as a valuable tool for studying neurodevelopmental disorders or nutritional challenges during this critical time-period.

**Figure 1.**
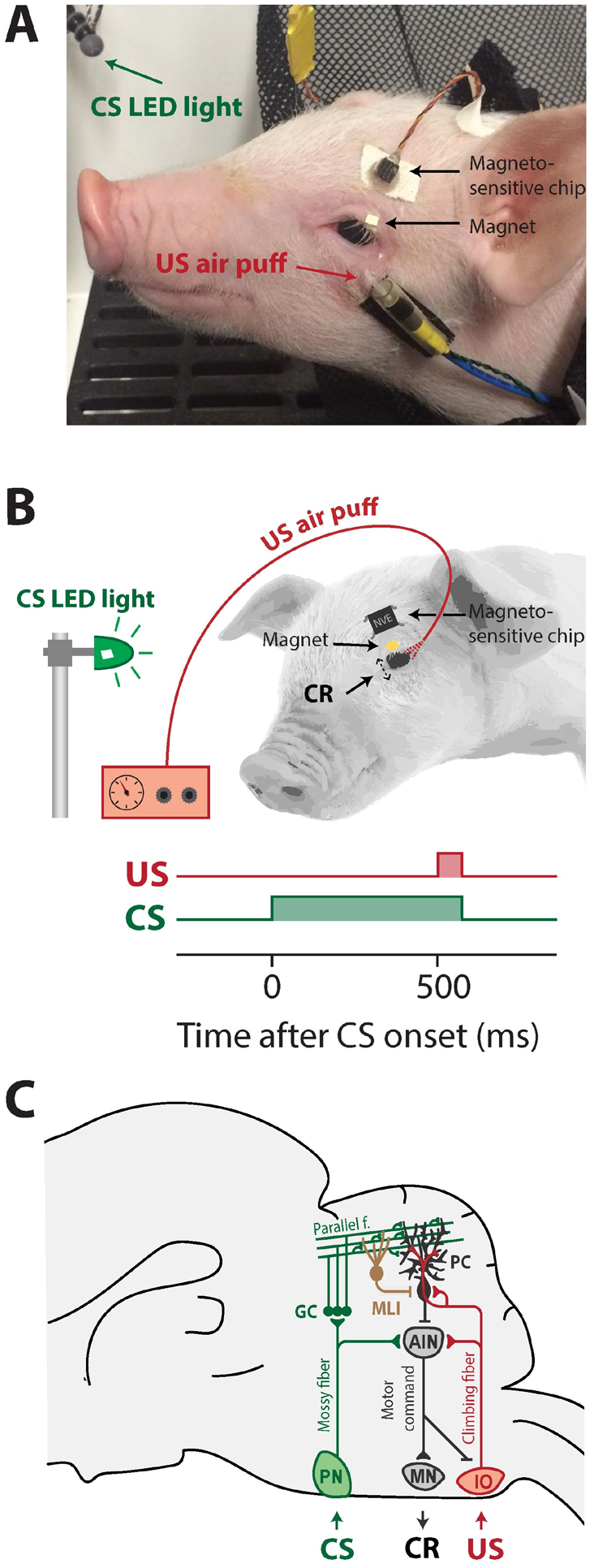
Overview of eyeblink conditioning apparatus, trial parameters, and neural circuitries involved in Pavlovian eyeblink conditioning. **(A)** Apparatus in place for eyeblink conditioning trials. The CS was a blue LED light, the US was a mild air-puff applied to the cornea. Since pigs tend to move their heads a lot during the experiment, the equipment needed to deliver the air-puff US and record the eyelid position were attached to the pig’s face, while the pig remained comfortably in a hammock. **(B)** Simplified illustration of the apparatus attachments and trial parameters of CS (green) and US (red). We used a delay eyeblink conditioning paradigm, whereby the start of the CS preceded the onset of the US by 500 ms and both CS and US co-terminated at 550 ms after CS onset. **(C)** Simplified illustration of the neural circuits underlying delay eyeblink conditioning. The CS is transmitted via the mossy fiber - parallel fibers system (green) to Purkinje cells in well-defined microzones in the cerebellar cortex. The same Purkinje cells receive input from the climbing fibers transmitting the US signal (red). Memory formation takes place at this point of convergence of the CS and US in the cerebellar cortex. CRs are driven by the cerebellar output to brainstem motor neurons that innervate the eyelid musculature. Abbreviations: AIN, anterior interposed nucleus; CR, conditioned response; CS, conditional stimulus; IO, inferior olive; MLI, molecular layer interneuron; MN, brainstem motor neurons innervating the eyelid musculature (N.III, N. VI, N. VII); PN, pontine nuclei; UR, unconditioned response; US, unconditional stimulus.

## 2. Materials and Methods

### 2.1 Animal housing and care

All animal care and experimental procedures were in compliance with the National Research Council Guide for the Care and Use of Laboratory Animal Care and Use Committee and approved by the University of Illinois Urbana-Champaign Institutional Animal Care and Use Committee. Beginning on postnatal day (PND) 2, eighteen naturally farrowed intact male pigs (n=18) that were the offspring of Line 2 boars and Line 3 sows (A Large White and Landrace cross, also known as “Camborough”. Pig Improvement Company, Henderson, TN) were artificially reared to PND 30 and provided a nutritionally complete milk replacer formula (Purina ProNurse, Land O’Lakes Inc., Arden Hills, MN, US). Pigs were individually housed in a custom artificial rearing system, which allowed the pigs to see, hear, and smell, but not touch the near-by pigs to closely control the individual cage environment (Mudd et al., 2016). Pigs were fed *ad libitum* using an automated milk replacer delivery system that dispensed milk from 1000 h to 0600 h the next day. Lights were automatically turned on at 0800 h and turned off at 2000 h. Daily observations and pig body weights were recorded to track clinical indicators (e.g., diarrhea, lethargy, weight loss, or vomiting).

### 2.2 Eyeblink conditioning system

. Each eyeblink conditioning experimental setup consisted of a solid, sound-damping box with a custom-designed hammock securely attached, in which the pig rested during trials. A blue light-emitting diode (LED) was attached inside the testing unit, approximately 10 cm from the anticipated location of the pig’s head. This blue LED light was used as a conditional stimulus (CS). Additionally, an airline connected to a regulator was attached to the pig’s head, just below the eye, and it was used as an unconditional stimulus (US). These pieces were attached using medical-grade surgical tape and glue **(Figure 1A)**. Individual blinks were determined by measuring the distance between a giant magnetoresistance (GMR) magnetometer (NVE, Eden Prairie, MN) adhered to the pig’s forehead and a small magnet adhered to the pig’s eyelid (Koekkoek et al., 2002). The magnetic distance measurement technique (MDMT) was the best option for detecting the eyelid position in pigs, since pigs tended to move their heads during the experiment which imposed serious constraints on the quality of high-speed video recordings. National Instruments (NI-PXI; Austin, TX) equipment was used to control experimental parameters and to acquire the eyelid position signal.

### 2.3 Trial parameters

The overall behavioral experiment consisted of two phases: habituation and acquisition. The habituation phase started at PND 14 and lasted for 5 consecutive days. During the habituation phase, the pigs acclimated to resting in the hammock and the testing environment without receiving any stimuli for 15 min each day. Following the habituation phase, the acquisition phase started at PND 19 as lasted for 5 consecutive days. Knowing that different species show different learning speeds (Rasmussen, 2019), we expected pigs to reach asymptotic levels of conditioned responses in 150-200 trials within 4-5 days. During the acquisition phase, the pigs were subjected to 40 trials per day for 5 consecutive days. There were a total of 5 blocks of trials per day, and each block contained 1 US-only trial, 6 paired trials, and 1 CS-only trial. The first 500 ms of each trial was a baseline period, followed by the onset of CS **(Figure 1B)**. The onset of US was 1000 ms after the beginning of the trial with 500 ms inter-stimulus interval (ISI), and US and CS co-terminated at 1050 ms after 50 ms of temporal overlap of CS and US. Inter-trial interval was determined by the following criteria: (1) a random duration between 8-12 sec at the minimum must pass; (2) the eye must at least be half open [i.e., fraction eyelid closure (FEC) must be equal or larger than 0.5]; and (3) the eyelid must be stable for at least 2 sec. A trained observer carefully monitored the pig and experimental parameters and adjusted equipment if necessary. The overall behavioral experiment started around the same time of the day, and a single handler stayed consistent for the entire experiment to minimize human handling bias.

### 2.4 Analysis of eyeblink conditioning data

Individual eyeblink traces were analyzed with custom computer software and R (v. 4.0.4, R Core Team, Austria). For each type of trial (i.e., CS-only, US-only, CS-US paired), a single snippet was taken from the MDMT eyelid position signal. Each snippet, hereafter called an ‘eyeblink trace’, had a duration of 2000 ms. Eyeblink traces were filtered in forward and reverse direction with a low-pass butterworth filter using a cutoff frequency at 50 Hz. Trials with significant activity in the 500 ms pre-CS period [> 7-times the interquartile range (IQR)] were regarded as invalid and disregarded for further analysis. Trials were normalized by aligning the 500 ms pre-CS baselines and normalizing the signal so that the size of a full blink was 1 FEC. This normalization was achieved by using the reflexive blinks to the air-puff (unconditioned responses, UR) as a reference.

For each session, we calculated the maximum value in the median eyelid trace plus one IQR and data were normalized by dividing each trace by this value. As a consequence, in the normalized traces, an FEC of 1 corresponded with the eye being fully closed, a FEC of 0 corresponded with the eye being fully open. In valid normalized CS-only and CS-US paired trials, all eyelid movements larger than 0.2 and with a latency to CR onset between 50-500 ms and a latency to CR peak between 150-500 ms (both relative to CS onset) were considered a CR. Additionally, we determined for each individual trial the following parameters: (1) FEC full interval: the maximum eyelid closure (= fraction eyelid closure) in the CS-US interval calculated over all valid trials wherein a CS was presented (CS-only and CS-US paired trials); (2) FEC at 500 ms: the maximum eyelid closure (= fraction eyelid closure) at the moment that the US is delivered (i.e., 500 ms after CS onset), calculated over all valid trials wherein a CS was presented (CS-only and CS-US paired trials); (3) FEC CR trials: the maximum eyelid closure (= fraction eyelid closure) in the CS-US interval calculated over the trials wherein a CR was present; (4) the latency to CR onset in trials wherein a CR was present; (5) the latency to CR peak in trials wherein a CR was present.

Because pigs responded with a partial eye opening to the CS at the start of training, we quantified for the amplitude (or strength) of the eyelid closures in response to the CS using three different outcome measures: (1) maximum amplitude of the eyelid closure in 150-500 ms interval after CS onset calculated over *all* trials (FEC_150-500_); (2) amplitude of the eyelid closure at 500 ms after CS onset calculated over *all* trials (FEC_500_); and (3) maximum amplitude of the eyelid closure in the 150-500 ms interval after the onset of the CS calculated over *only the trials wherein a CR was present* (CRamplitude_150-500_).

Statistical analysis was done using multilevel linear mixed-effects (LME) models in R Studio (code available upon request). The LME have several major advantages over standard parametric and non-parametric tests (Aarts et al., 2014; Schielzeth et al., 2020), as they are more robust to violations of normality assumptions, which is often the case in biological data samples. Moreover, use of LME models are able to accommodate the nested structure of our data (i.e., trial nested within session, session nested within animal, animal nested within group). Finally, LME models are objectively better at handling missing data points than repeated measures analysis of variance (ANOVA) models and do not require homoscedasticity as an inherent assumption. In our LME, we used session as a fixed effect, and pig as a random effect. (R code: lme (outcome ~ session_nr), control = ctrl, data = df1, random = ~ session_nr | pig_id, method = “REML”, na.action=na.exclude), Covariance structure: unstructured). Goodness of fit model comparison was determined by evaluating log likelihood ratio, BIC, and AIC values. The distribution of residuals was inspected visually by plotting the quantiles of standard normal versus standardized residuals (i.e., Q-Q plots). Data were considered as statistically significant if the p-value was smaller than 0.05.

## 3. Results

All pigs were trained for five consecutive days in the delay Pavlovian eyeblink conditioning test. The training protocol used the association of a 550-ms LED light flash as the CS, ending with a 50-ms air-puff delivered to the pig’s cornea as the US, which triggered the involuntary eyeblink response (i.e., UR).

### 3.1 Eye openings in response to the novel LED serving as conditional stimulus

When inspecting the raw eyeblink traces, we noticed that pigs often responded with a further opening of the eye (represented by a decrease below zero in fraction eyelid closure) in response to the CS during the first two training sessions (**Figure 2A, B, Figure 3A)**. Examination of the MDMT signal and videos during the eyeblink conditioning test suggested that pigs often had the upper eyelid partly closed, whereby the upper eyelid was not naturally in the fully opened position. This partial eye closure was the neutral position of the pig’s eye and we confirmed that it was not artificially produced by instrumentation around the eye. As a consequence, the averaged eyeblink traces (**Figure 3A**) for testing days 1 and 2 show a clear eye opening in response to the CS (see Discussion).

**Figure 2.**
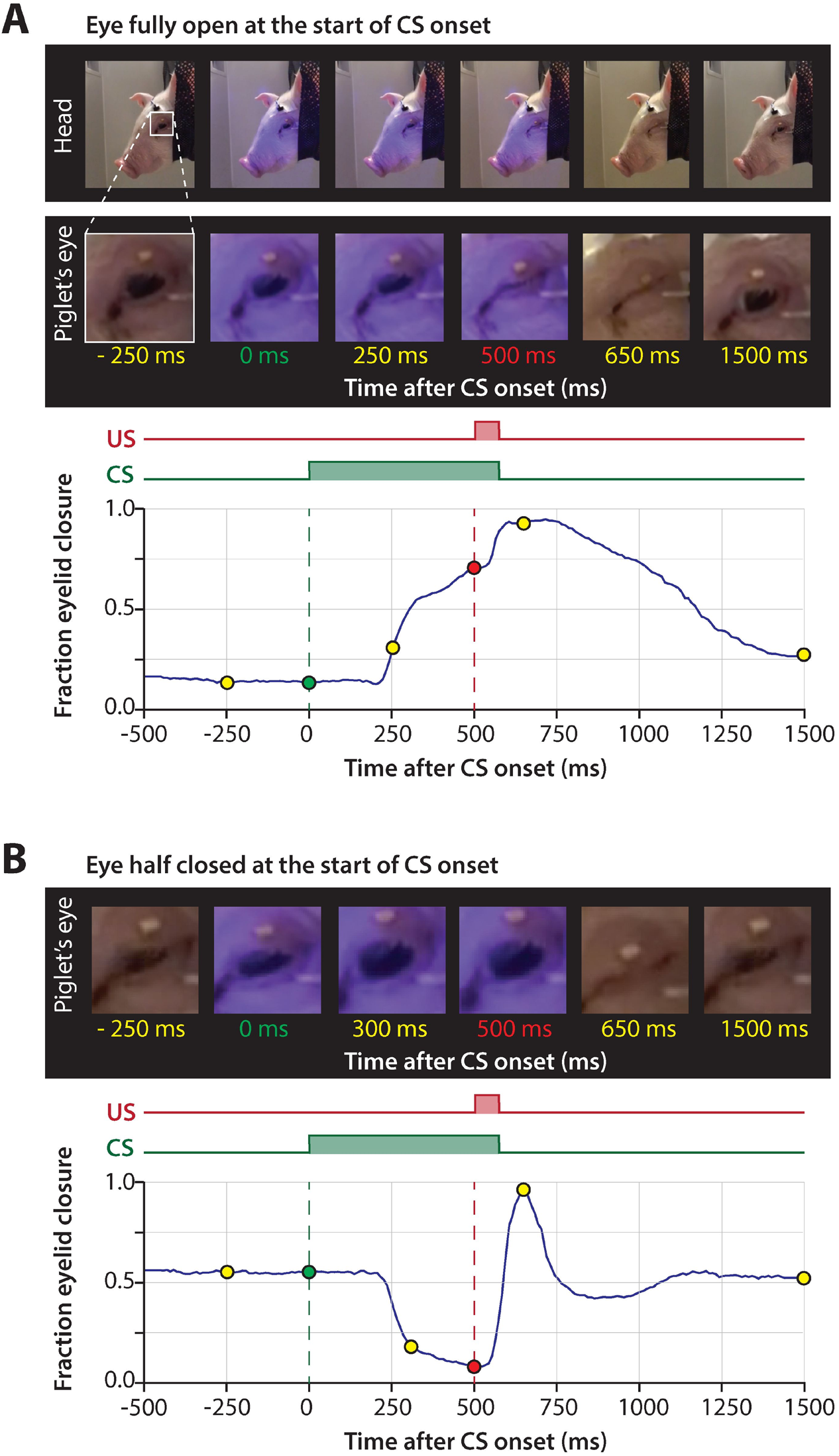
Representative eyelid responses to the CS. **(A)** The eyelid is fully opened at the start of CS onset during the first two training sessions, demonstrated by a series of pictures at different time-points and corresponding eyelid trace. The blue LED used as a CS is reflected on the skin of the pig. **(B)** Similar to but now the eyelid is half-closed at the start of CS onset during the last training session. Abbreviations: CS, conditional stimulus; US, unconditional stimulus.

**Figure 3.**
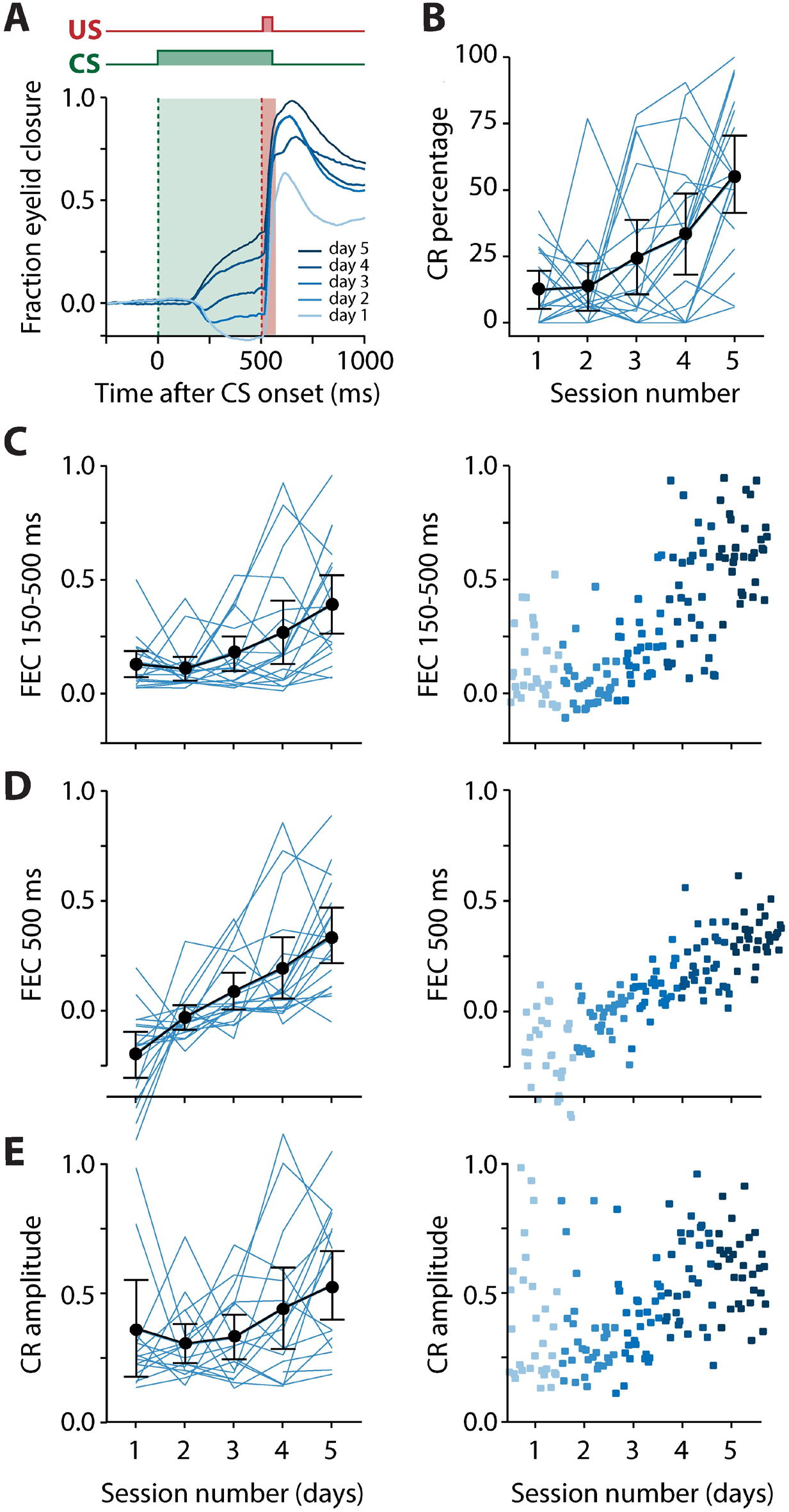
Pigs learn the eyeblink conditioning task, shown by a gradual increase of the CR probability and amplitude of eyelid responses to the CS. **(A)** Averaged eyeblink traces for each training session. A fraction eyelid closure (FEC) of 0 corresponds with full eye opening, while an FEC of 1 corresponds with a full eyelid closure. During the first and second training sessions, the averaged eyeblink traces exhibit a prominent eye opening in response to the CS. Prolonged training (sessions 3-5) resulted in eyelid closures in response to the CS, which are considered as conditioned responses (CR). **(B)** Individual pig learning curves (thin light blue lines) and group-averaged learning curve (thick black line) showing the CR probability as a function training session. Error bars indicate the 95% confidence interval. A statistically significant effect of session was found for CR probability. **(C-E)** Left panels: individual pig learning curves (thin light blue lines) and group-averaged learning curve (thick black line) as a function of session. Error bars indicate the 95% confidence interval. Right panels: trial-by-trial values whereby each dot represents the average for all values for that trial. The colors (ranging from light blue to dark blue) correspond with the colors of the averaged eyeblink traces in panel A. **(C)** The maximum amplitude of the eyelid closure in 150-500 ms interval after CS onset calculated over *all* trials (FEC_150-500_). A statistically significant effect of session was found for FEC_150-500_. **(D)** The amplitude of the eyelid closure at 500 ms after CS onset calculated over *all* trials (FEC_500_) A statistically significant effect of session was found for FEC_500_. **(E)** The maximum amplitude of the eyelid closure in 150-500 ms interval after the onset of the CS calculated over *only the trials wherein a CR was present* (CRamplitude_150-500_). A statistically significant effect of session was found for CRamplitude_150-500_. For all statistical effects, please refer to **Table 1**. Abbreviations: CR, conditioned response; CS, conditional stimulus; FEC, fraction eyelid closure.

### 3.2 CR probability and the amplitude of eyelid responses to the CS

We found a significant main effect of day on CR probability (F(4,68)=11.75, p<0.0001, ANOVA on LME) **(Figure 3B, Table 1)**. A CR was defined as an FEC larger than 0.2 (full closure is 1, full opening is 0) in the interval of 150 to 500 ms after CS onset. On average, pigs started with a probability of 12.37 [±7.02 95% confidence interval (CI)] on session 1 and reached a CR probability of 55.42 (±14.39 95% CI) on day 5 **(Figure 3B, Table 1)**. The averaged FEC_150-500_ showed a statistically significant effect of session (F(4,68) = 6.05, p = 0.0003, ANOVA on LME). On average, pigs started with a FEC_150-500_ of 0.12 (±0.06 95% CI) on session 1 and reached a value of 0.39 (±0.12 95% CI) on day 5 **(Figure 3C, Table 1)**. Similarly, for FEC_500_ there was a main effect of day (F(4,48)=11.79, p<0.0001, ANOVA on LME). On average, pigs started with a FEC500ms of −0.19 (±0.10 95% CI) on day 1 and reached a value of 0.33 (±0.12 95% CI) at the end of training **(Figure 3D, Table 1).** Note the negative value on day 1 for FEC_500_, reflecting the partial eyelid opening, that is not masked when only looking at the FEC_150-500_. Finally, we also found a small but significant main effect of session for CRamplitude_150-500_ (F(4,61) = 3.04, p = 0.02). On average, pigs started with a CRamplitude_150-500_ of 0.33 (±0.17 95% CI) on day 1 (although there were only a handful of CRs on day 1, most of them probably being spontaneous blinks that were indistinguishable from learned CRs) and ended with an CRamplitude_150-500_ of 0.48 (±0.12 95% CI) **(Figure 3E, Table 1).**

**Table 1.** Average percentage, amplitude, and timing of conditioned responses in all trials^1^

| Session<sup>2</sup> | CR Percentage<br>(0.2 FEC criterion) | Max FEC between 150 and<br>500 ms after CS onset | FEC at 500 ms<br>after CS onset | CR amplitude | Latency to CR<br>onset | Latency to CR<br>peak | SD Latency to<br>CR peak |
| --- | --- | --- | --- | --- | --- | --- | --- |
| 1 | 12.37 ($\pm 7.02$) | 0.12 ($\pm 0.06$) | -0.19 ($\pm 0.10$) | 0.33 ($\pm 0.17$) | 102.52 ($\pm 43.15$) | 299.16 ($\pm 61.47$) | 131.55 ($\pm 27.39$) |
| 2 | 13.39 ($\pm 8.81$) | 0.11 ($\pm 0.06$) | -0.03 ($\pm 0.05$) | 0.27 ($\pm 0.07$) | 100.91 ($\pm 34.29$) | 347.02 ($\pm 59.66$) | 103.23 ($\pm 42.60$) |
| 3 | 24.59 ($\pm 13.87$) | 0.17 ($\pm 0.08$) | 0.08 ($\pm 0.08$) | 0.29 ($\pm 0.07$) | 118.95 ($\pm 33.74$) | 394.45 ($\pm 40.71$) | 78.40 ($\pm 23.24$) |
| 4 | 33.15 ($\pm 15.07$) | 0.27 ($\pm 0.14$) | 0.19 ($\pm 0.13$) | 0.40 ($\pm 0.14$) | 116.56 ($\pm 28.84$) | 401.62 ($\pm 32.15$) | 98.88 ($\pm 25.97$) |
| 5 | 55.42 ($\pm 14.39$) | 0.39 ($\pm 0.12$) | 0.33 ($\pm 0.12$) | 0.48 ($\pm 0.12$) | 146.85 ($\pm 29.75$) | 420.74 ($\pm 22.67$) | 78.59 ($\pm 17.21$) |
| Session<sup>3</sup> | F(4,68) = 11.75,<br>p < 0.0001 | F(4,68) = 6.05, p = 0.0003 | F(4,48) = 11.79,<br>p < 0.0001 | F(4,61) = 3.04,<br>p = 0.02 | F(4,61) = 1.49,<br>p = 0.21 | F(4,61) = 3.86,<br>p = 0.0007 | F(4,54) = 2.91,<br>p = 0.03 |

<sup>1</sup>Abbreviation: CR, conditioned response; FEC, fraction eyelid closure; SD, standard deviation.<sup>2</sup>All values from sessions 1 to 5 are mean ± 95% CI (confidence interval).<sup>3</sup>All F-values and p-values are derived from ANOVA on linear mixed-effects model.

### 3.3 Adaptive timing of the conditioned responses

Next, we examined more closely the adaptive timing of the eyeblink CR over the course of training. For CR timing, we quantified three different outcomes only in trials wherein a CR was present: (1) latency in milliseconds to the onset of eyelid CR in the interval between 150 and 500 ms after CS onset; (2) latency in milliseconds to the maximum peak of the CR in the interval between 150 and 500 ms after CS onset. Note that the onset of the air-puff US is at 500 ms after CS onset. In addition, we looked at the variability of the latencies to CR peak, since we observed in the raw traces that over the course of training the timing of these CR peaks became more precise, i.e., became more centered around the onset of the air-puff US.

The latency to CR peak time lacked a temporal preference in the first session, but clearly showed preference from the moment that animals started to show CRs more reliably in session 2 and onwards. The averaged latency to CR peak showed a statistically significant effect of session (F(4,61) = 3.86, p = 0.007, ANOVA on LME). On average, pigs started with a latency to CR peak of 299.16 (±61.47 95% CI) ms on session 1 and a value of 420.74 (±22.67 95% CI) ms on the last day of training **(Figure 4A, C, Table 1),** and herewith the eyelid was maximally closed exactly at the moment that the US would be delivered. Because we noted that variability became smaller over time, we also quantified the standard deviation of latency to CR peak for each session and each pig. The averaged standard deviation of CR peak time latencies showed a statistically significant effect of session (F(4,54) = 2.91, p = 0.03, ANOVA on LME). On day 1 we calculated an average standard deviation of 131.55 (±27.39 95% CI) milliseconds, and this value got gradually smaller, reaching a minimum value of 78.59 (±17.21 95% CI) on day 5 (**Figure 4B, Table 1)**. The averaged latency to CR onset showed no statistically significant effect of session (F(4,61) = 1.49, p = 0.21, ANOVA on LME). On average, pigs started with a latency to CR onset of 102.52 ms (±43.15 95% CI) on session 1 and this value stayed relatively stable with a value 146.85 (±29.75 95% CI) ms on session 5 **(Figure 4D, Table 1)**. Based on these timing parameters of the eyeblink CR, we conclude that pigs were able to adaptively time their eyeblink CR.

**Figure 4.**
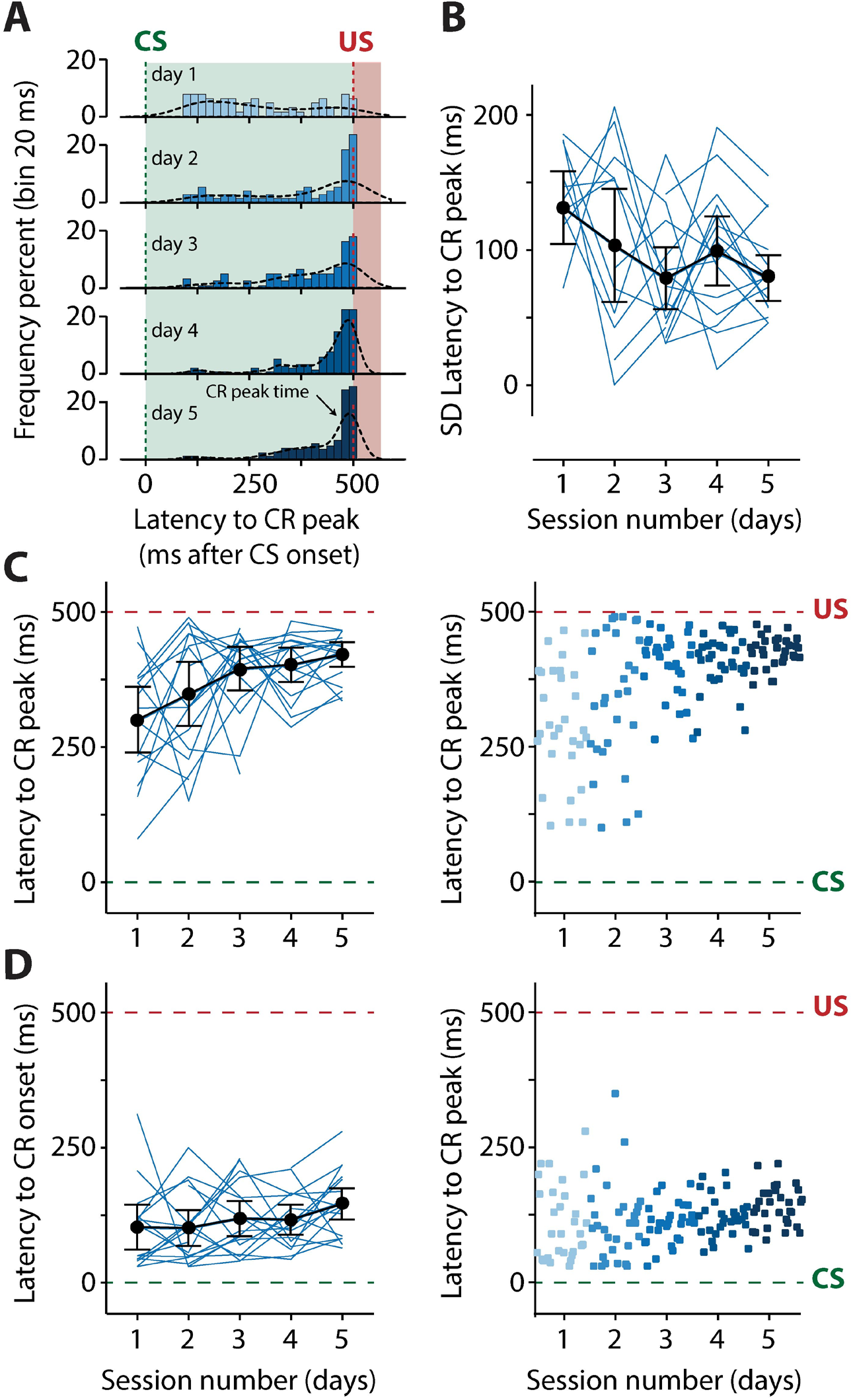
Pigs learn the eyeblink conditioning task, as shown by the timing of the conditioned eyelid responses. **(A)** Development of the distribution of the latency to CR peak from day 1 (top panel) through day 5 (bottom panel). The green dashed line indicates CS onset and the red dashed line indicates US onset. The latency to CR peak time lacked a temporal preference in the first session, but clearly showed preference from the moment that animals started to show CR more reliably in session 2 and onwards. The latency to CR peak was very close to the moment that the US would be delivered. **(B)** Standard deviation of the latency to CR peak as a function of session. Thin light blue lines represent individual pig curves and the thick black line is the group-averaged curve. Error bars indicate the 95% confidence interval. A statistically significant effect found of session was found for the SD of the latency to CR peak, indicating that the variability in the CR peak times get smaller over time **(C, D)** Left panels: individual pig learning curves (thin light blue lines) and group-averaged learning curve (thick black line) as a function of session. Error bars indicate the 95% confidence interval. Right panels: trial-by-trial values whereby each dot represents the average of all animals for all values for that trial. The colors (ranging from light blue to dark blue) correspond with the colors of the averaged eyeblink traces in panel A. Green dashed line indicates CS onset, red dashed line indicates US onset. **(C)** The latency in milliseconds to the maximum peak of the CR in the interval between 150 and 500 ms after CS onset. Note how the CR peaks concentrate around the onset of the US. A statistically significant effect found of session was found for the latency to CR peak. **(D)** The latency in milliseconds to the onset of eyelid CR in the interval between 150 and 500 ms after CS onset. The latency to CR onset remained stable over time. No statistically significant effect found of session was found for the latency to CR onset. Abbreviations: CR, conditioned response; CS, conditional stimulus; SD, standard deviation, US, unconditional stimulus.

## 4. Discussion

The main purpose of the present study was to develop and validate the eyeblink conditioning paradigm in young pigs. We found that pigs were indeed able to learn the eyeblink conditioning task: both CR probability and the CR amplitude showed a gradual increase over the course of five days of training. Moreover, the eyeblink CR were properly timed, in the sense that the eyelid was maximally closed exactly around the onset of the air-puff US, herewith providing the optimal protection against the aversive air-puff while perturbing the pig’s vision for the shortest amount of time. Thus, we show for the first time that eyeblink conditioning can reliably be performed in young pigs, herewith providing a new neurobehavioral task to study the effect of nutrition on cerebellar development in pigs.

### 4.1 Learning rates and CS-US interval

It is known that different species learn at different rates. Humans often need only a single session of 50 paired CS-US trials to learn the task (Knickmeyer et al., 2008; Thürling et al., 2015), for rabbits it often takes about 5-6 days of about 50 paired CS-US trials (Gormezano et al., 1962; Welsh & Harvey, 1989), and mice and rats are closer to 8-9 days of 100 paired CS-US trials a day to reach asymptotic levels of conditioning (Albergaria et al., 2018). Based on these findings, we estimated that it would require a pig to reach asymptotic levels of conditioning in 4-5 days. However, our data shows that five days of conditioning with 40 paired CS-US trials per day was not sufficient to reach these asymptotic levels. Future studies on pig eyeblink conditioning should consider using either more trials per day or more days of training, to reach higher values for CR probability and CR amplitude.

The duration of the CS-US interval has an effect on learning speed. Mice learn the delay eyeblink task the quickest at intervals close to 200 ms (Chettih et al., 2011; Heiney et al., 2014). For humans and rabbits, intervals around 500 ms are commonly used and induce reliable conditioning. For that reason, we also chose an interval of 500 ms between CS and US onset for the pigs, and it appeared that pigs learned reasonably well in this interval. It fell beyond the scope of this study to extensively investigate what the optimal interval is for pigs to learn the eyeblink conditioning task, so further investigation in this area is warranted.

### 4.2 Eye openings in response to the novel conditional stimulus

As mentioned above, we observed that pigs often responded with a further opening of the eye in response to the CS during the first two training sessions (**Figure 2A, B, Figure 3A)**. Examination of the MDMT signal and videos during the eyeblink conditioning test taught us that pigs often had the eyelid partly closed, whereby the upper eyelid was dropped down a bit. This partial eye closure was the neutral position of the pig’s eye and was not due to any instrumentation around the eye, since we observed the same eyelid position during the habituation sessions when there was not equipment attached to the pig’s face and even when the animals were just in their home cage. As a consequence, the averaged eyeblink traces (**Figure 3A**) for days 1 and 2 show a clear eye opening in response to the CS. These eye openings were not considered a CR, since they were even present from the start of training and even during the habituation sessions before any CS-US pairings had occurred. Whereas the language may be construed as speculative, we considered this phenomenon as a sign of the pig’s curiosity to the novel stimulus. A similar, but much more subtle, response has been reported in mice (Grasselli et al., 2020). Prolonged training of the pigs led to eyelid closures instead of openings in response to the CS, which reinforces the plastic nature of the response. These eyeblink closures, as seen on sessions 3, 4, and 5, were considered as true eyeblink CR (**Figure 3A**).

### 4.3 Pigs as an appropriate model for studies on nutrition and brain development

Translating infancy to young pigs, one month in pig’s life equates to roughly one year in human’s life in terms of their total brain volume growth (Thibault & Margulies, 1998). Thus, our 4-to-5-wk-old pigs can be considered as 4-to-5-month-old infants. Interestingly, infants aged four to five months, or even younger, have shown their capability of performing delay eyeblink conditioning (Lintz et al., 1967; Ivkovich et al., 1999; Herbert et al., 2003). During this critical period, delay eyeblink conditioning can serve as a sensitive behavioral paradigm to study cerebellar development.

As briefly discussed above, the pigs have similar nutrient requirements and intestinal anatomy and physiology to those of humans (Odle et al., 2014), and these similarities allow the pigs to be utilized as an established preclinical and translational model, especially in nutritional neuroscience studies. Nutrition in neonatal pigs has profound and wide-ranging effects on neurodevelopment. Regarding cerebellar development, it has been demonstrated using Magnetic resonance imaging that an iron deficiency in pigs results in a decrease in relative cerebellum volume as they aged from PND 32 to 61, suggesting iron deficiency results in neurodevelopmental alterations of the cerebellum (Mudd et al., 2018).

While eyeblink conditioning has been suggested as a valuable biomarker for defining several neurological disorders such as fetal alcohol syndrome and autism spectrum disorder (Reeb-Sutherland & Fox, 2013), eyeblink conditioning has not been extensively utilized in nutritional developmental neuroscience. In this regard, we consider studies involving the direct effects of caffeine or alcohol on eyeblink conditioning performance as separate from nutritional intervention, and thus, believe they are not particularly useful for nutritional and developmental neuroscience. Only one study involving the effects of perinatal iron deficiency in rats employed eyeblink conditioning, where the nutrient deficiency elicited mild to severe impairments in eyeblink conditioning performance (McEchron et al., 2008). Thus, as nutritional deficiency and supplementation during early-life can influence brain development and cognitive functions (Mudd et al., 2016; Liu et al., 2014; Fleming et al., 2019), sensitive behavioral paradigms such as eyeblink conditioning in pigs can serve as valuable methodological tools to investigate how these dietary changes can influence cognitive and behavioral development.

### 4.4 MDMT to detect eyelid movement

The field of eyeblink conditioning is moving away from electromyography and MDMT towards high-speed video recordings to detect eye blinks. We started our eyeblink experiments in pigs with high-speed video recordings but noticed that the pigs tend to move their heads a lot during experimentation, making it hard for a single camera in the eyeblink chamber to capture the eye. Modern, smaller cameras that are attached to the pig’s head could potentially solve this problem. Still, the MDMT was able to detect eyelid movements at high spatiotemporal resolution while putting minimal restraint to the pigs by allowing relatively free head movement.

### 4.5 Quiet wakefulness

One of the noteworthy technical challenges with eyeblink conditioning in young pigs is controlling their alertness level. We observed pigs sometimes displayed squinting behaviors during the inter-trial intervals, and some pigs were even falling asleep. Incidents of a loss of alertness and falling asleep especially occurred in the later part of the training (sessions 3, 4, 5). This is rather challenging to work with, as it may result in reduced reactions to the CS and US, resulting in decreased CR performance levels; therefore, the level of alertness can greatly influence the performance in eyeblink conditioning. This behavior is not exclusively present in young pigs, but it is a technical challenge that must be considered. It closely resembles the state of quiet wakefulness that has been reported for mice when the eyeblink conditioned task was performed in the home cage of the animal (Boele et al., 2010). The state of quiet wakefulness in mice is described as “the animal sitting quietly sitting in the corner of its cage with its eyes partially closed”, and it is not thought to be an anxiety-related freezing behavior (Boele et al., 2010). In mice, it appeared that making the eyeblink task more engaging, by letting the animal walk on a treadmill system for instance, was more successful than modifications such as mild food restrictions or delivery of short auditory cues to startle the animals. Similarly, in pigs one might consider ways to make the eyeblink task more engaging, for instance by introducing a screen to watch a movie or the placement of new objects for the animal to explore during the task.

## 5. Conclusion

As use of the pig as a preclinical and translational model in nutritional neuroscience research has been gaining its popularity, development of sensitive behavioral paradigm for pigs has been critical to accurately measure changes in cognitive functions that may have been resulted from nutritional changes. Pigs that are 4 weeks of age are capable of performing eyeblink conditioning, as demonstrated by the CR percentage, the amplitude of the eyelid responses to the CS, and the improvement in the timing of their CR across sessions. However, five days of training with 40 paired trials per session does not appear to be sufficient to observe asymptotic learning. Overall, the current experiment was the first study to demonstrate that eyeblink conditioning in four-week-old pigs may serve as a sensitive and valuable behavioral paradigm to measure cognitive development.

## Author Contributions

Conceptualization: HJB, SJ, AM, SF, SK, RND

Methodology: HJB, AM, SF, SK, JF, RND

Investigation: HJB, SJ, AM, SF, SK, JF

Analysis: HJB, AM

Visualization: HJB

Funding acquisition: RND

Project administration: HJB, SJ, RND

Supervision: SK, RND

Writing – original draft: HJB, SJ

Writing – review & editing: All authors

## Abbreviations

CS: Conditional stimulus
US: unconditional stimulus
CR: conditioned response
PND: postnatal day
MDMT: magnetic distance measurement technique
ISI: inter-stimulus interval
FEC: fraction eyelid closure
UR: unconditioned response
LME: linear mixed-effects models

## Acknowledgements

The authors would like to thank Kristen Karkiewicz and Adam Jones for daily animal care.

